# The genetic architecture of structural left-right asymmetry of the human brain

**DOI:** 10.1101/2020.06.30.179721

**Authors:** Zhiqiang Sha, Dick Schijven, Amaia Carrion-Castillo, Marc Joliot, Bernard Mazoyer, Simon E. Fisher, Fabrice Crivello, Clyde Francks

## Abstract

Left-right hemispheric asymmetry is an important aspect of healthy brain organization for many functions including language, and can be altered in cognitive and psychiatric disorders^1-8^. No mechanism has yet been identified for establishing the human brain’s left-right axis^9^. We performed multivariate genome-wide association scanning (mvGWAS) of cortical regional surface area and thickness asymmetries, and subcortical volume asymmetries, using data from 32,256 participants from the UK Biobank. There were 21 significant loci affecting different aspects of brain asymmetry, with functional enrichment involving microtubule-related genes and embryonic brain expression. These findings are consistent with a known role of the cytoskeleton in left-right axis determination in other organs of invertebrates and frogs^10-12^. Genetic variants affecting brain asymmetry overlapped with those influencing autism, educational attainment and schizophrenia.

---

Only three loci have previously been reported at a genome-wide significant level to affect brain asymmetries, for specific features of temporal lobe anatomy^13,14^, and without revealing the broader biological pathways involved. For each of the 32,256 participants, and each of 73 bilaterally paired regional measures of brain structure, we calculated hemispheric asymmetry indexes, AI=(left-right)/(left+right)/2 (for 33 cortical surface area AIs, 33 cortical thickness AIs, and 7 subcortical volume AIs (Supplementary Table 1). The measures were derived from cortical parcellation and subcortical segmentation of T1 brain images (Methods). All but one of the regional mean AIs were significantly different from zero, indicating population-level asymmetries (Bonferroni corrected p<0.05, Supplementary Fig. 1 and Supplementary Table 2), in left-right directions consistent with previous reports^5,6^. For example, some language-related regions showed greater average left than right surface areas, including superior temporal and supramarginal cortex, and pars opercularis. Language is an archetypal lateralized function, for which most people have left-hemisphere dominance^15^.

Forty-two AIs showed significant single-nucleotide-polymorphism-based (SNP-based) heritabilities (FDR-corrected p<0.05), ranging from 2.2% to 9.4%, i.e. 28 of the surface area AIs, 8 cortical thickness AIs, and 6 subcortical volume AIs (Fig. 1A, Supplementary Table 3). The overall pattern was consistent with previous twin-based heritability analyses^5,6^. SNP-based genetic correlation analysis indicated overlapping genetic contributions to some of the AIs (Fig. 1B, Supplementary Fig. 2, Supplementary Tables 4-10). Within some cortical regions, surface area and thickness AIs had negative genetic correlations (Supplementary Table 10), which indicates that variants can have antagonistic effects on surface and thickness asymmetries of these regions.

**Figure 1.**
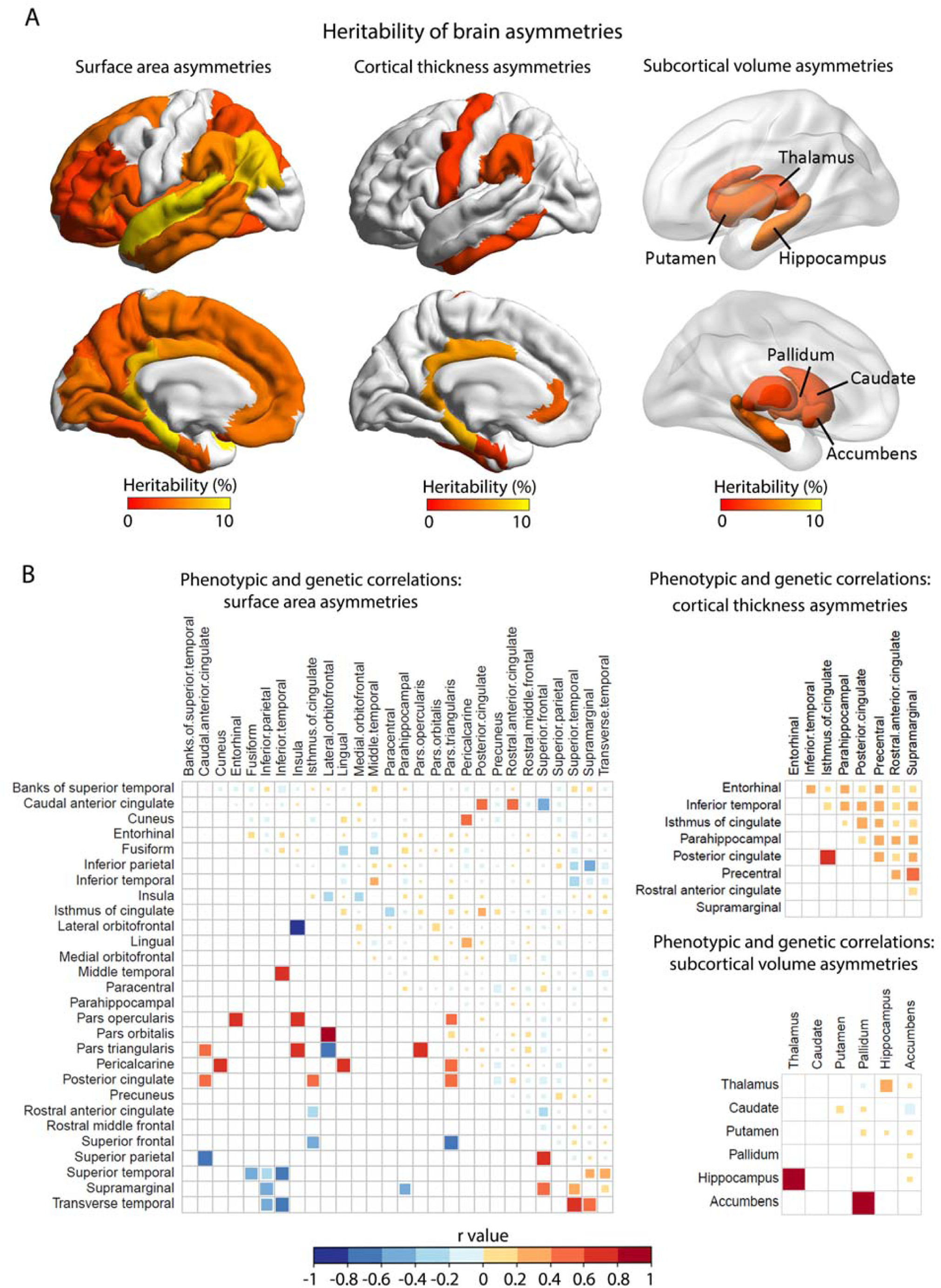
SNP-based heritability and correlation analysis of regional brain asymmetry measures. (A) SNP-based heritability estimates for brain asymmetry measures. Only regions for which AIs were significantly heritable are indicated in color. (B) Genetic and phenotypic correlations between AIs. Phenotypic and genetic correlations between each pair of AIs are in the upper right and lower left triangles, respectively. Only significantly heritable AIs are shown that also have at least one significant phenotypic or genetic correlation after FDR correction. The sizes and colours of the squares indicate the correlation coefficients.

We performed mvGWAS for 9,803,522 SNPs using meta-canonical correlation implemented in MetaPhat^16^, using the 42 AIs with significant SNP-based heritability. In this way, one mvGWAS was used to screen the genome for association simultaneously with 42 AIs. FUMA^17^ was used to clump mvGWAS results based on linkage disequilibrium (LD), and identify lead SNPs. There were 21 distinct genomic loci at the 5×10^−8^ significance level (Fig. 2, Table 1 and Supplementary Fig. 3), represented by 27 independent lead SNPs (with pairwise LD r^2^<0.1) (Table 1).

**Table 1.**
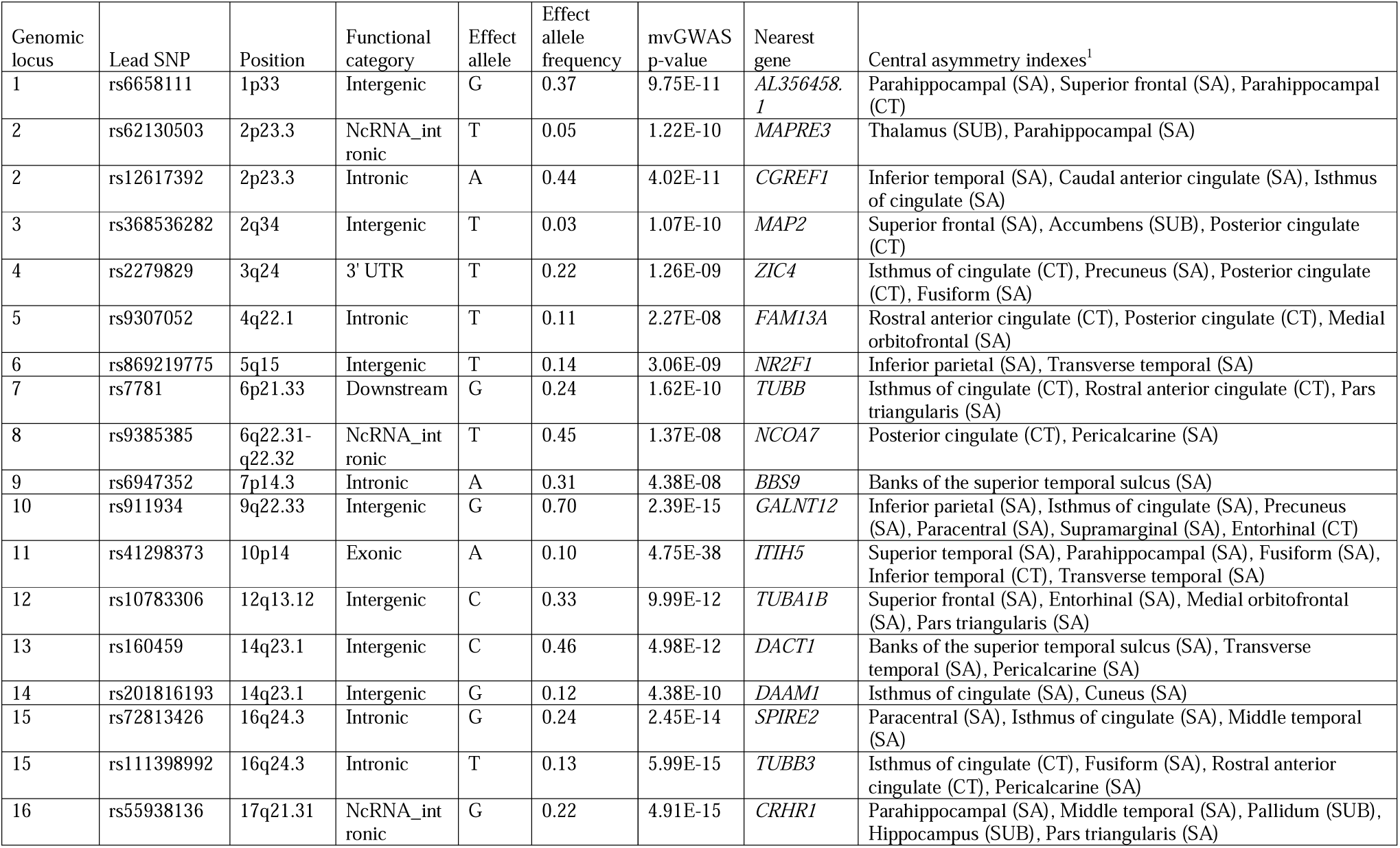
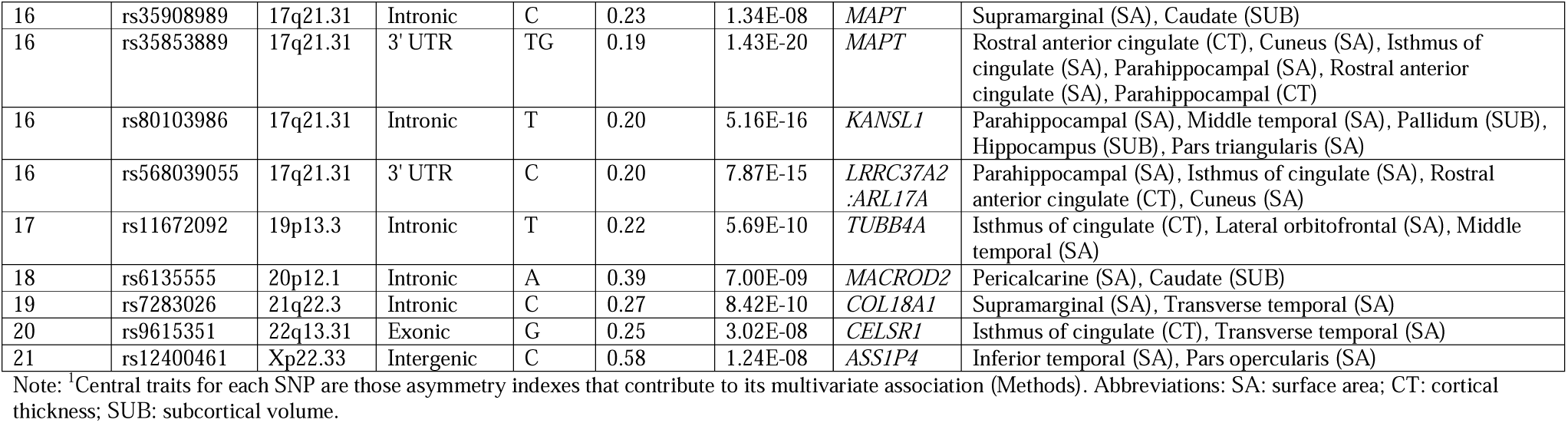
Genomic loci affecting brain asymmetries. All lead SNPs are shown.

**Figure 2.**
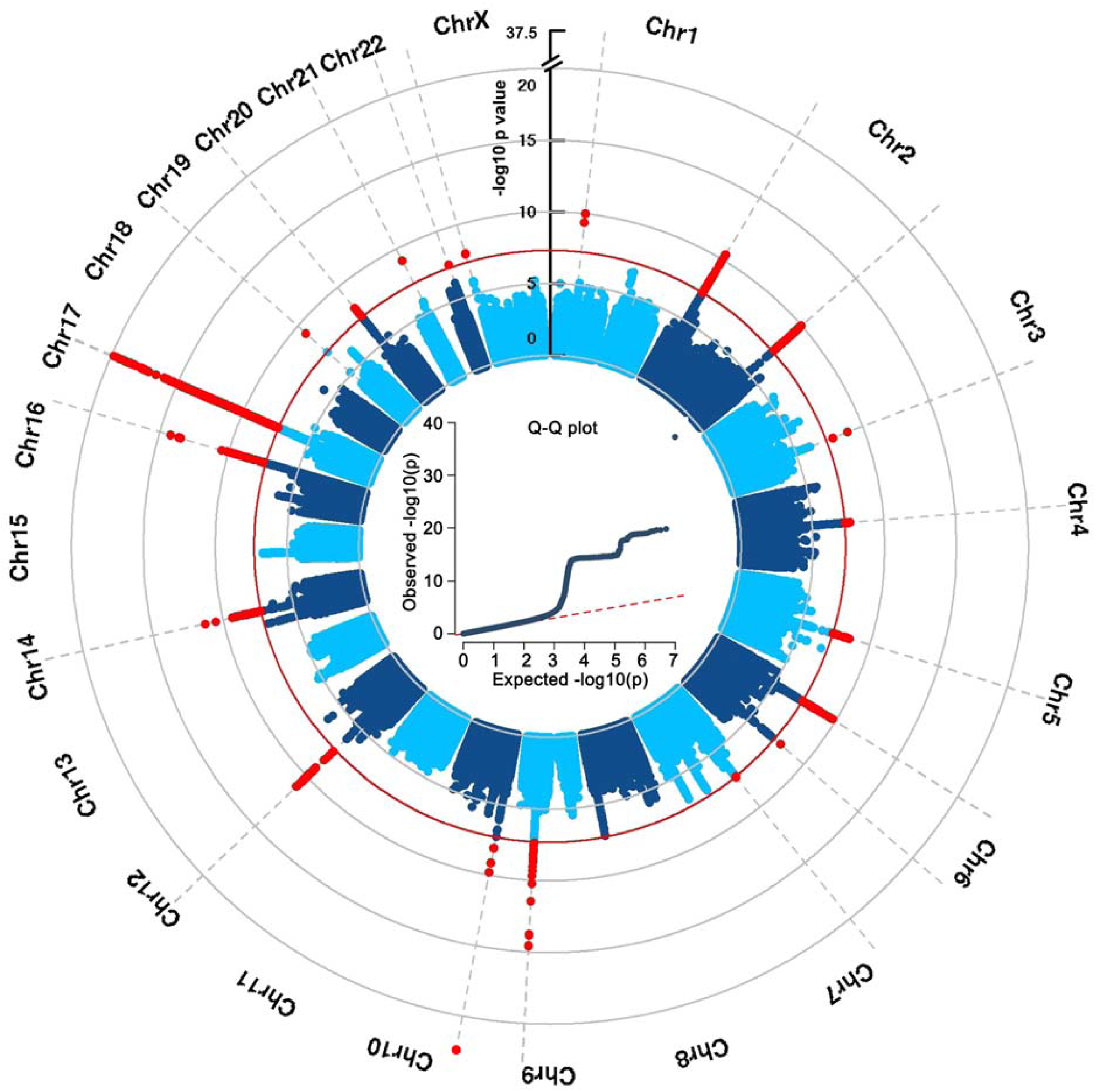
Multivariate GWAS analysis of regional brain asymmetries in 32,256 participants. Circular Manhattan plot for multivariate GWAS across asymmetries of surface area, cortical thickness and subcortical volumes. The red dashed line indicates the significance threshold p<5×10^−8^ (Methods). The Q-Q plot is shown at the center.

For each lead SNP, phenotype decomposition^16^ identified the ‘central’ AIs that contributed to its multivariate association (Supplementary Table 11). Most central AIs affected by the 27 lead SNPs were distributed in core regions of the language (e.g. lateral temporal, pars opercularis, supramarginal) and limbic systems (e.g. cingulate, orbitofrontal and mesial temporal cortex; Fig. 3). For example, the most significant SNP, with p=4.75×10^−38^ (rs41298373 on 10p14) had five central AIs: the minor allele was associated with a leftward shift of surface area asymmetry for two lateral temporal regions, a rightward shift of surface area asymmetry for two medial temporal regions, and a leftward shift of cortical thickness asymmetry in the inferior temporal gyrus (Supplementary Table 11). A locus on 17q21 affected asymmetries of cortical surface area, thickness and subcortical volume (Table 1). The effects of lead variants separately on left and right hemispheric measures are shown in Supplementary Table 11.

**Figure 3.**
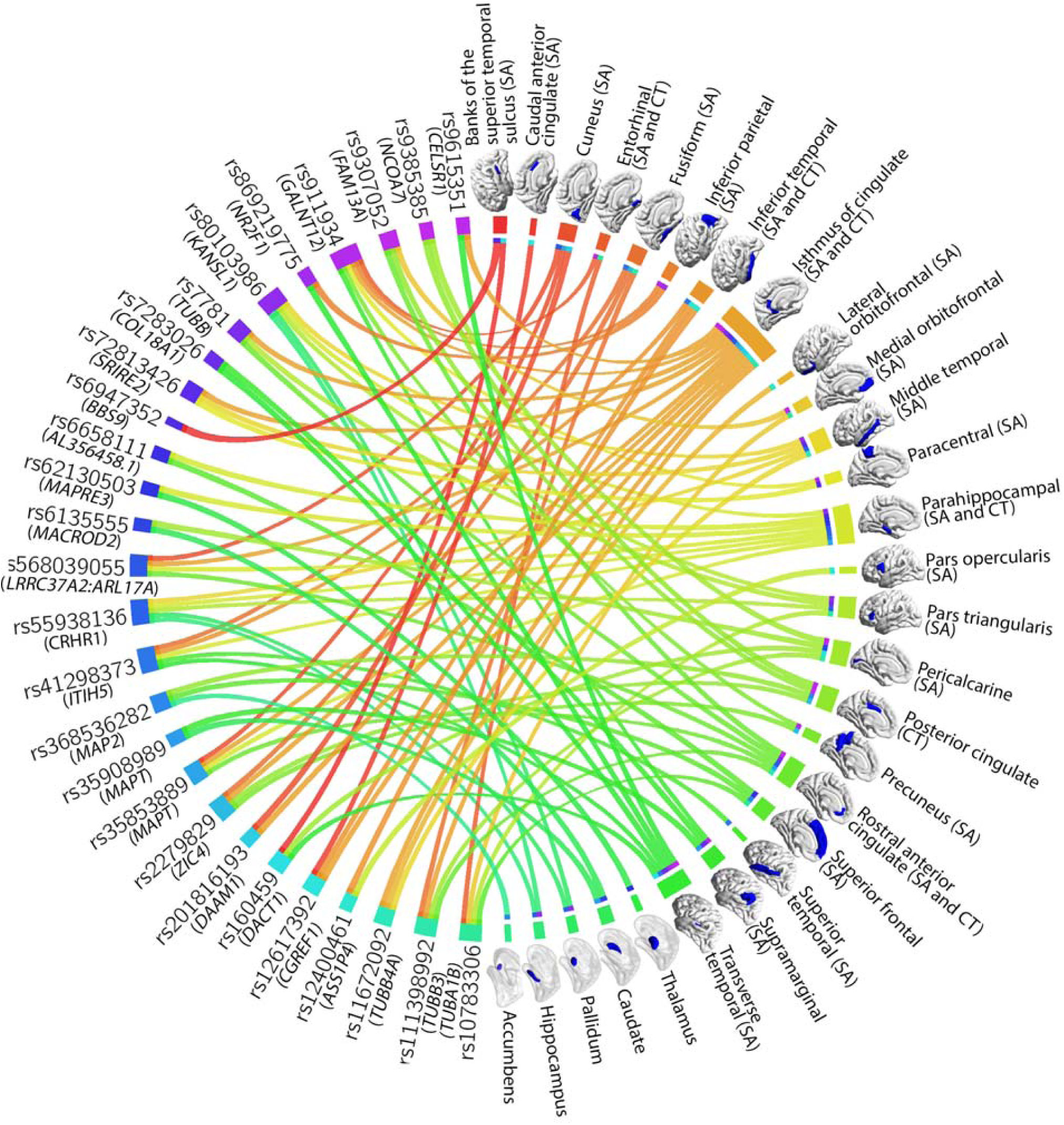
Overview of 27 independent lead variants associated with different regional brain asymmetries. Circle plot illustrating the 27 lead variants from mvGWAS (left) in relation to the central asymmetry indexes (right) underlying their specific multivariate associations. Different colors indicate different lead variants or regional asymmetries. The closest genes to the lead variants are shown. It can be seen that most central asymmetry indexes are of regional surface areas, and that some variants affected multiple asymmetries of different types. Abbreviations: SA: surface area; CT: cortical thickness; SUB: subcortical volume.

FUMA^17^ applied three strategies to annotate candidate SNPs to genes at significantly associated loci (Methods): physical position, expression quantitative trait locus (eQTL) information, and chromatin interactions (Supplementary Table 12 and Supplementary Figs 4-5). Here we summarize notable annotations for the lead SNPs: On 1p33, rs6658111 is close to *AL356458*.*1*, a pseudogene of *MTMR14* (myotubularin related protein 14). On 2p23.3, rs62130503 is intronic to *MAPRE3* (microtubule associated protein RP/EB family member 3a), and rs12617392 is a brain eQTL^18^ of *MAPRE3*. On 2q34, rs368536282 is close to *MAP2* (microtubule associated protein 2), a well-known dendrite-specific marker of neurons^19^, previously implicated in left-handedness by GWAS analysis^20-22^. On 3q24, rs2279829 is a cortical eQTL^18^ of *ZIC4*, which is involved in visual and auditory pathway development^23^. rs9307052 on 4q22.1 is in high LD (r^2^=0.99) with the handedness-associated variant rs28658282^21^. On 5q15, rs869219775 is close to *NR2F1*, which is involved in neural activity during cortical patterning^24^. On 6p21.33, rs7781 is in the 3’ untranslated region (UTR) of *TUBB* (tubulin beta class I). On 7p14.3, rs6947352 is intronic to *BBS9*, which causes Bardet-Biedl syndrome when mutated, involving retinopathy and intellectual disability^25,26^. On 9q22.33, rs911934 is located in a region having a chromatin interaction with *TRIM14* in adult cortex^27^ (Supplementary Fig. 4), a gene which may activate Wnt/β-catenin signaling and affects mesodermal versus ectodermal differentiation of embryonic stem cells^28^. On 10p14, rs41298373 is a predicted deleterious missense coding variant in *ITIH5*, which was previously reported to affect *planum temporale* volumetric asymmetry^14^. ITI family proteins are involved in extracellular matrix stabilization^29^. On 12q13.12, rs10783306 is close to the alpha tubulin gene *TUBA1B*. This variant is also in high LD with a handedness-associated variant, rs11168884 (r^2^=0.89)^21^. On 14q23.1, two lead variants for two independent genomic loci, rs160459 and rs201816193, showed evidence for cross-locus chromatin interaction via the promoters of nearby genes in fetal cortex^27^ (Supplementary Fig. 4). The former is near to *DACT1*, a locus which has been reported to affect superior temporal sulcus depth^13^, while the latter is close to *DAAM1*, which modulates the reorganization of the actin cytoskeleton and the stabilization of microtubules^30,31^. Two lead variants on 16q24.3, rs72813426 and rs111398992, are in introns of *SPIRE2* and the tubulin gene *TUBB3* respectively, both of which are key proteins in cytoskeleton organization^32,33^. On 17q21.31 there were five independent lead SNPs: rs35908989 is intronic to *MAPT* which encodes microtubule-associated protein tau, and rs55938136, rs35853889 and rs568039055 are brain eQTLs^18,34,35^ of *MAPT*, while rs80103986 is in high LD (r^2^=0.91) with handedness-associated variant rs55974014^21^. On 19p13.3, rs11672092 is intronic to the tubulin gene *TUBB4A*, and in high LD with rs66479618 (r^2^=0.88), another handedness-associated SNP^21^. On 20p12.1, rs6135555 is in a region having a chromatin interaction with the *FLRT3* promoter in neural progenitor cells^36^ (Supplementary Fig. 4), a gene which regulates axon guidance and excitatory synapse development^37^. On 21q22.3, rs7283026 is intronic to *COL18A1*, involved in neural tube closure and mutated in Knobloch syndrome^38^, which can include skull abnormalities. On 22q13.31, rs9615351 is an exonic variant of a gene involved in planar cell polarity, *CELSR1*^39^. On Xp22.33, rs12400461 is close to pseudogene *ASS1P4* and upstream of *MXRA5*; the latter encodes a matrix remodeling-associated protein and is implicated in autism^40^.

To further link asymmetry-associated variants to genes, genome-wide gene-based association analysis^41^ was performed based on the results from mvGWAS. There were 112 significant genes at Bonferroni-corrected P< 0.05 (Supplementary Fig. 6 and Supplementary Table 13). Seven of these genes were previously associated with handedness^21^: *MAP2, FAM13A, TUBA1B, TUBB3, CRHR1, RABAC1 and TUBB4A*. Seventy-two of the 112 genes have been reported to associate with educational attainment^42^ and 16 with intelligence^43^ (Supplementary Table 14). Sixty-two of the 112 genes were also mapped by at least one of the three SNP-to-gene mapping strategies above (Supplementary Table 13). Of these 62 genes, there were 51 with proteins annotated in the STRING database^44^, among which there were 117 known or putative interactions, compared to 8 interactions expected for a random set of this size from the whole proteome (p<1×10^−16^). Microtubule-related genes (e.g. *MAP2, MAPT, SPIRE2* and *TUBA1A*) were centered at the hubs and connectors of the largest network (Fig. 4A). We also used the genome-wide, gene-based p values for enrichment analysis using MAGMA^41^, in relation to 7,343 Gene Ontology ‘biological process’ sets defined within MSigDB^45^. The gene sets ‘regulation_of_microtubule_binding’ (p=1.52×10^−6^) and ‘negative_regulation_of_neuron_differentiation’ (p=5.59×10^−6^) showed significant enrichment (adjusted P<0.05, Bonferroni correction, Supplementary Table 15). Significant enrichment within various microtubule-related sets was also found when using the list of single closest genes (Table 1) to the 27 lead SNPs (Supplementary Table 16). Enrichment in microtubule-related sets was not reported in a recent GWAS of bilaterally averaged cortical surface area and thickness measures in 51,665 individuals^46^, suggesting a particular involvement in hemispheric asymmetry rather than bilateral measures.

**Figure 4.**
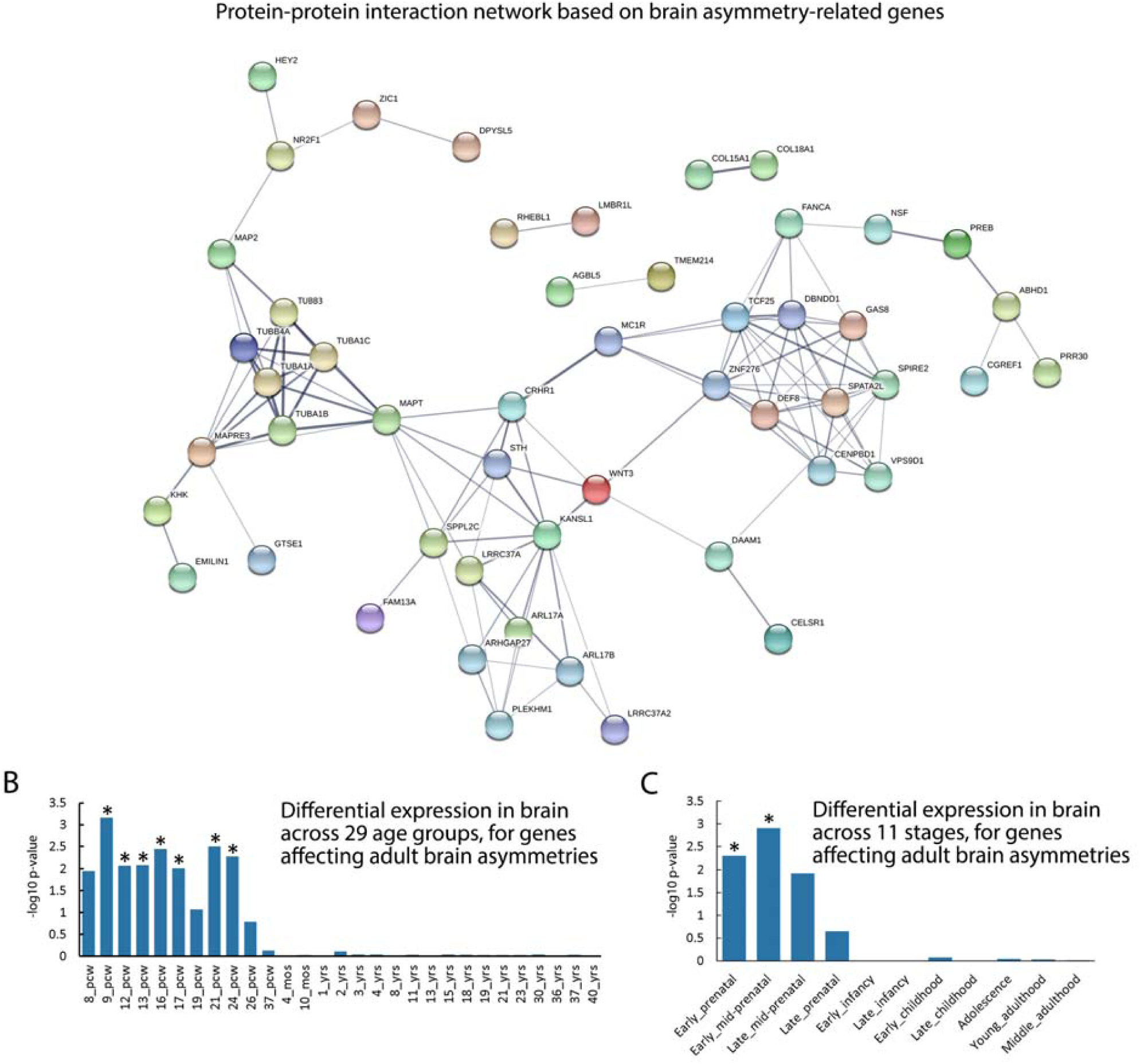
Functional annotations of variants affecting brain asymmetry. (A) Asymmetry-associated genes integrated into a protein-protein interaction network. Proteins are represented by nodes. Edges between nodes represent protein-protein associations and are weighted by confidence scores provided in the STRING database (Methods). Colored nodes represent the queried proteins. Only medium-confidence (>0.4) links were retained, and disconnected proteins are not shown. (B) Relation between gene-based association with brain asymmetries and higher mRNA expression in the human brain at particular ages, using BrainSpan data from 29 age groups. Asterisks represent the significant age groups meeting FDR correction with p<0.05. (C) Relation between gene-based association with brain asymmetries and higher mRNA expression in the human brain at particular ages, using BrainSpan data from 11 defined age groups. Asterisks represent the significant groups meeting FDR correction with p<0.05.

Testing our genome-wide, gene-based P values with respect to BrainSpan^47^ human gene expression data from either 29 age groups, or 11 defined developmental stages, we found higher mRNA expression of brain-asymmetry-associated genes during early-prenatal (p=5.17×10^−3^) and early-mid-prenatal (p=1.25×10^−3^) stages, from 9 (p=6.95×10^−4^) to 24 (p=8.62×10^−3^) post-conceptional weeks (FDR corrected p values <0.05) (Figs. 4B and 4C, Supplementary Table 17). This is consistent with the fact that various anatomical asymmetries of the brain are already visible *in utero*^48,49^, and supports the existence of an early developmental mechanism for establishing the brain’s left-right axis^50-52^.

We next used iSECA to perform genetic overlap analyses^53^ with our mvGWAS results in relation to GWAS summary statistics from neurodevelopmental disorders, behavioral and psychological traits which have been reported to associate phenotypically with aspects of structural and/or functional brain asymmetry: attention deficit hyperactivity disorder^54-58^, autism spectrum disorder^59-64^, educational attainment^42,65,66^, handedness^4,22,67^, intelligence^43,68-71^ and schizophrenia^72-77^. There was evidence for genetic overlap between brain asymmetries and autism (p=0.005), educational attainment (p=0.001) and schizophrenia (p=0.002) which remained significant at Bonferroni-corrected P<0.05 (Fig. 5, Supplementary Figs. 7-8 and Supplementary Table 18). Further research will be needed to understand whether brain asymmetries mediate gene-trait associations in a causal sense. Although we did not observe genetic overlap of brain asymmetry with handedness at a genome-wide level, we did note individual loci in common between these traits (above). In addition, we found no overlap between our mvGWAS results and those from a previous GWAS of intracranial volume in 32,438 participants^78^, (Supplementary Table 18, Supplementary Figs. 7-8), which again indicates that the genetic architecture of brain asymmetry is largely distinct from brain size.

**Figure 5.**
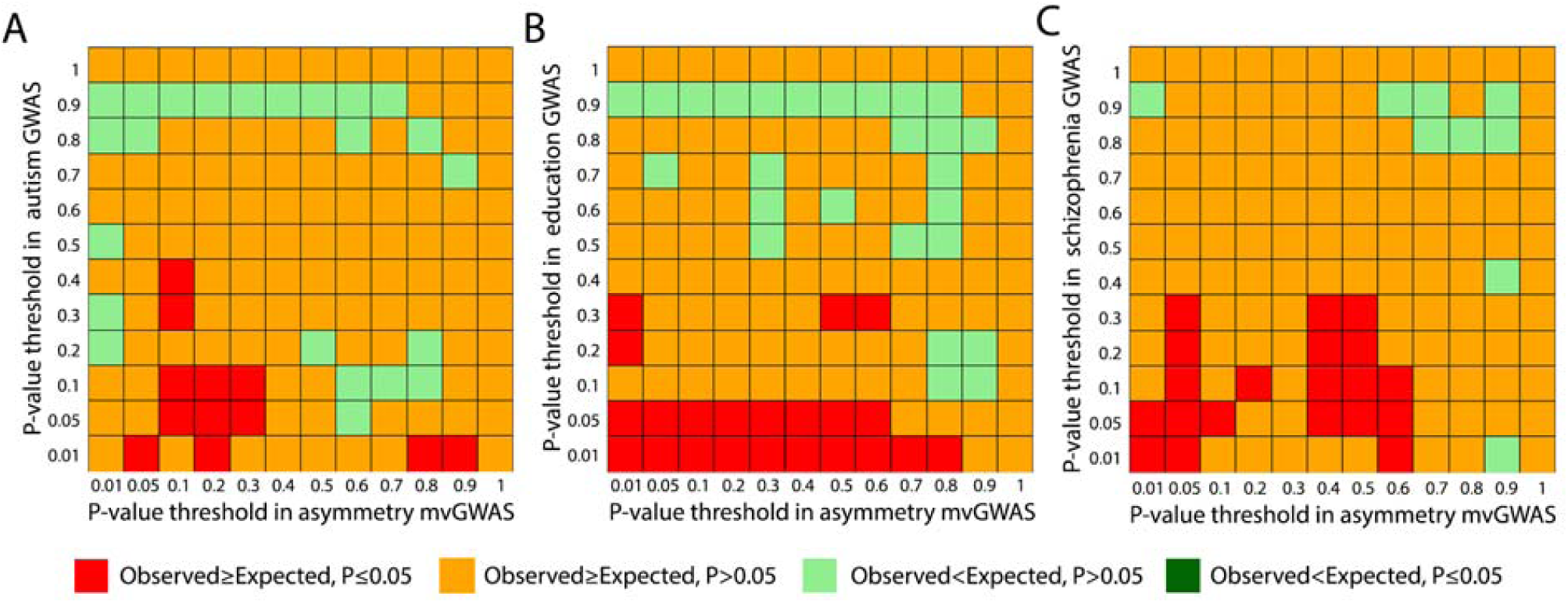
Genetic overlaps between brain asymmetries and other traits. Heatmap plots illustrating pleiotropic effects between brain asymmetries and autism (A), educational attainment (B) and schizophrenia (C), based on per-SNP genome-wide association scan (GWAS) p values for these traits from previous studies (Methods), in relation to the multivariate GWAS (mvGWAS) p values from the present study of brain asymmetries.

Previous studies in invertebrates and frog embryos have shown that the cytoskeleton plays a role in cellular chirality and the establishment of the left-right axis of other organs, in a non-ciliary-dependent manner^10,12,79,80^. Our findings motivate genetic-developmental studies of left-right differentiation of the embryonic mammalian brain, focused on a possible cytoskeletal-mediated mechanism of axis formation. Such a mechanism may be organ-intrinsic, i.e. distinct from other pathways that establish broader aspects of body asymmetry^10,79,81^. While many brain asymmetries were strong and directional at the population level, their heritabilities were low. This suggests that genetic-developmental mechanisms for brain asymmetry are tightly constrained and largely genetically invariant in the population, and that environmental factors and/or developmental randomness are responsible for most variability^22,82-84^. A cytoskeleton-based origin of brain asymmetry would fit this scenario, as the cytoskeleton is essential for fundamental functions in cellular biology, beyond axis formation^85,86^.

## Online Methods

### Participants

This study was conducted under UK Biobank application 16066, with Clyde Francks as principal investigator. This is a general adult population cohort. The UK Biobank received ethical approval from the National Research Ethics Service Committee North West-Haydock (reference 11/NW/0382), and all of their procedures were performed in accordance with the World Medical Association guidelines. Informed consent was obtained for all participants. We used the brain imaging data released in February 2020, and data availability and processing (discussed below) resulted in a final sample of 32,256 participants of white British ancestry, together with the structural magnetic resonance imaging data and genotype data from the same participants. The age range of these participants was from 45 to 81 years (mean 63.77), and 15,288 were male, 16,968 were female.

### Genetic quality control

We downloaded imputed SNP genotype data from the UK Biobank data portal (bgen files; imputed data v3-release March 2018). We first excluded subjects with a mismatch of their self-reported and genetically inferred sex, with putative aneuploidies, or who were outliers based on heterozygosity (principle component corrected heterozygosity>0.19) and genotype missingness (missing rate>0.05)^87^. All the analyses were restricted to participants with ‘British ancestry’, which was defined by Bycroft et al. (‘in.white.British.ancestry.subset’)^87^. We randomly excluded one subject from each pair with a kinship coefficient >0.0442, as defined within the UKB relatedness file ‘ukb1606_rel_s488366.dat’. Next, QCTOOL (v.2.0.6) and PLINK^88^ were used to perform genotype quality control: excluding SNPs with minor allele frequency <1%, Hardy-Weinberg equilibrium test p-value<1×10^−7^, and imputation INFO score <0.7 (a measure of genotype imputation confidence). We also excluded multi-allelic SNPs because most of the downstream software (below) could not handle them. This resulted in 9,803,522 bi-allelic variants.

### Neuroimaging phenotypes and covariates

Brain anatomical measures of regional cortical surface area, cortical thickness and subcortical volumes were derived from the T1 assessments (Siemens Skyra 3T MRI with 32-channel RF receive head coil) released by the UK Biobank Imaging Study (for the full protocol: http://biobank.ndph.ox.ac.uk/showcase/refer.cgi?id=2367). Briefly, *in vivo* whole brain T1-weighted MRI scans were used to perform cortical parcellation into 34 regions per hemisphere with the Desikan-Killiany Atlas^89^, and 7 subcortical structural segmentations. Surface area was measured at the grey-white matter boundary, and thickness was measured as the average distance in a region between the white matter and pial surfaces. Details of the image quality control and processing are described elsewhere^90^. Given that the data for the temporal pole were reported as unreliable^90^, we only used 33 surface area, 33 cortical thickness and 7 subcortical volume measures in each hemisphere (Supplementary Table 1). Per measure, we removed data-points greater than six standard deviations from the mean. Then, we calculated the AI for each matching pair of left and right measures, in each participant, as (left-right)/((left+right)/2). Given this definition, a positive AI reflects leftward asymmetry (greater left than right). The AI is a widely used measure in brain asymmetry studies^5,91,92^. The denominator ensures that the index does not simply scale with brain size, i.e. the left-right difference is adjusted for the bilateral measure. For each AI, one-sample t testing was used to examine whether the population mean AI was significantly different to zero, with Bonferroni correction at 0.05 for multiple testing. Subsequently, the distributions of AIs were normalized by rank-based inverse normalization to minimize statistical artifacts. The normalized AIs were used as input for subsequent analysis.

The Desikan-Killiany atlas^89^ was derived from manual segmentations of sets of reference brain images. The labelling system incorporates hemisphere-specific information on sulcal and gyral geometry with spatial information regarding the locations of brain structures, and shows a high accuracy when compared to manual labelling results^89^. Accordingly, the mean regional asymmetries in the UK Biobank might partly reflect left-right differences present in the reference dataset used to construct the atlas. However, our study was focused primarily on comparing *relative* asymmetry between genotypes, at the regional level. The use of an asymmetrical atlas based on healthy individuals had the advantage that regional identification was likely to be accurate for structures that are asymmetrical in the general population, while taking hemisphere-specific information into account.

We also made use of continuous variables as covariates in heritability estimation and genome-wide association analysis (below), which were: age when attended assessment center (UKB field IDs 21003-2.0), nonlinear age (zage^2^), first ten genetic principle components capturing population genetic diversity (UKB field ID: 22009-0.1∼22009-0.10), scanner position parameters (UKB field IDs of X, Y and Z position: 25756-2.0, 25757-2.0 and 25758-2.0), T1 signal-to noise ratio (UKB field ID: 25734-2.0) and T1 contrast-to-noise ratio (UKB field ID: 25735-2.0), plus categorical covariates which were: assessment center (UKB field ID: 54-2.0), genotype measurement batch (UKB field ID: 22000-0.0) and sex (UKB field ID: 31-0.0).

### SNP-based heritability and genetic correlation analysis within the UK Biobank data

Specifically for SNP-based heritability and genetic correlation analyses within the UK Biobank data, we further excluded one random participant from each pair having a kinship coefficient higher than 0.025 before genetic relatedness matrix construction (as this analysis is especially sensitive to higher levels of relatedness), resulting in 30,315 participants for these particular analyses. 9,516,074 autosomal variants with minor allele frequencies>1%, INFO score>0.7 and Hardy-Weinberg equilibrium p>1×10^−7^ were used to build a genetic relationship matrix using GCTA^93^ (version 1.93.0beta). Genome-based restricted maximum likelihood (GREML)^93^ analyses were performed to estimate the SNP-based heritability for each AI, controlling for the above-mentioned covariates, and applying FDR 0.05 across the 73 AIs to define significantly heritable AIs. Bivariate GREML^94^ analysis was used to estimate genetic correlations between pairs of AIs, separately for cortical surface area, cortical thickness and subcortical volume AIs, with FDR correction at 0.05 for multiple testing.

### Multivariate genome-wide association analysis

In mvGWAS, a single association test is performed for each SNP in relation to multiple traits simultaneously. We used MetaPhat^16^ to perform mvGWAS analysis across asymmetries for cortical surface area, cortical thickness and subcortical volume, including only the 42 AIs that had shown significant SNP-based heritability. MetaPhat performs meta-canonical correlation analysis, and uses univariate GWAS summary statistics as input from each separate AI, which were derived under an additive genetic model while controlling for the above-mentioned covariates, using BGENIE (v1.2)^87^. Thus our mvGWAS tested effectively for association with 42 traits. This approach estimates the linear combination of traits that is maximally associated with genotype, which can differ for each SNP, while maintaining a correct false positive rate. 9,803,522 SNPs (see further above) were used for mvGWAS, spanning all autosomes and chromosome X. Statistically significant SNPs were considered as those with P<5×10^−8^ in mvGWAS, which is a widely used threshold to account for multiple testing over the whole genome, in the context of LD in European-descent populations^95,96^.

MetaPhat also uses systematic criteria to define central traits which make the greatest contributions to significant multivariate associations, based on an iterative process to optimize multivariate model properties with reference to canonical correlation analysis p-values and the Bayesian Information Criterion^16^. For the lead SNPs at genome-wide significant loci (see below for how these were defined), we also performed post-hoc analysis in which we examined their separate left and right hemispheric effects, using traits corresponding to the central AIs that were involved in the multivariate associations (Supplementary Table 11), again using BGENIE, an additive genetic model, and covariates as described above.

As a sensitivity analysis, we re-ran the mvGWAS after excluding 886 participants who had lifetime diagnoses of neurological conditions that could potentially disrupt brain structure: ICD9 or ICD10 neurological diagnoses defined in Chapter I “Certain infectious and parasitic diseases.”, Chapter II “Neoplasms”, Chapter V “Mental and behavioral disorders.”, Chapter VI “Diseases of the nervous system”, Chapter IX “Diseases of the circulatory system.”, Chapter XVI “Certain conditions originating in the perinatal period.” and Chapter XVII “Congenital malformations, deformations and chromosomal abnormalities.” (Supplementary Table 19). The significant mvGWAS loci were minimally affected by this exclusion (Supplementary Fig. 9).

### Identification of genomic risk loci and functional annotations

FUMA (version v1.3.5e)^17^, an online platform for functional annotation of GWAS results, was applied to the results from mvGWAS. A multi-step process, using default parameters, was used to identify distinct, significantly associated genomic loci, and independent lead SNPs within those loci. Briefly, based on pre-calculated LD structure from the 1000G European reference panel^97^, SNPs with genome-wide significant mvGWAS P values <5×10^−8^ that had LD r^2^<0.6 with any others were identified. For each of these SNPs, other SNPs that have r^2^≥0.6 with them were included for further annotation (see below), and independent lead SNPs were also defined among them as having low LD (r^2^<0.1) with any others. If LD blocks of significant SNPs are located within 250□kb of each other (default parameter), they are merged into one genomic locus. Therefore, some genomic loci could include one or more independent lead SNPs (Table 1). The major histocompatibility complex region on chromosome 6 was excluded from this process by default, due to its especially complex and long-range LD structure.

Functional annotations were applied by matching chromosome location, base-pair position, reference and alternate alleles to databases containing known functional annotations, which were ANNOVAR^98^ categories, Combined Annotation-Dependent Depletion^99^ scores, RegulomeDB^100^ scores, and chromatin state^101,102^:

1. ANNOVAR categories identify SNPs based on their locations with respect to genes, such as exonic, intronic and intergenic, using Ensembl gene definitions.
2. Combined Annotation-Dependent Depletion scores predict deleteriousness, with scores higher than 12.37 suggesting potential pathogenicity^103^.
3. RegulomeDB scores integrate regulatory information from eQTL and chromatin marks, and range from 1a to 7, with lower scores representing more importance for regulatory function (see Supplementary Fig. 10 legend).
4. Chromatin states show the accessibility of genomic regions, and were labeled by 15 categorical states (see Supplementary Fig. 10 legend) based on 5 chromatin marks for 127 epigenomes in the Roadmap Epigenomics Project^102^, which were H3K4me3, H3K4me1, H3K36me3, H3K27me3 and H3K9me3. For each SNP, FUMA calculated the minimum chromatin state across 127 tissue/cell-type in the Roadmap Epigenomics Project^102^. Categories 1-7 are considered as open chromatin states.

We also used FUMA to annotate independent significant SNPs and their candidate SNPs according to previously reported phenotype associations (p<5×10^−5^) in the NHGRI-EBI catalog^104^.

For a significant mvGWAS association in the major histocompatibility complex region (Table 1), we took the most significant individual SNP, rs7781 (p=1.62×10^−10^), as the single lead SNP to represent this locus, and annotated it manually.

### SNP-to-gene mapping

SNP-to-gene mapping at significant mvGWAS loci was performed using the default FUMA processes for these three strategies:

1. Positional mapping was used to map SNPs to protein-coding genes based on physical distance (within 10kb) in the human reference assembly (GRCh37/hg19).
2. eQTL mapping was used to annotate SNPs to genes (i.e. when SNP genotypes are associated with variation in gene mRNA expression levels). eQTL mapping was carried out in relation to genes up to 1Mb away based on four brain-expression data repositories: PsychENCORE^35^, CommonMind Consortium^18^, BRAINEAC^34^, GTEx v8 Brain^105^. FUMA applied a FDR of 0.05 within each analysis to identify significant eQTL associations.
3. Chromatin interaction mapping was performed to map SNPs to genes based on seven brain-related Hi-C chromatin conformation capture datasets: PsychENCORE EP link (one way)^35^, PsychENCORE promoter anchored loops^18^, HiC adult cortex^27^, HiC fetal cortex^27^, HiC (GSE87112) dorsolateral prefrontal cortex^36^, HiC (GSE87112) hippocampus^36^ and HiC (GSE87112) neural progenitor cell^36^. We further selected only those genes for which one or both regions involved in the chromatin interaction overlapped with a predicted enhancer or promoter region (250 bp up- and 500 bp downstream of the transcription start site) in any of the brain-related repositories from the Roadmap Epigenomics Project^102^, i.e. E053 (neurospheres) cortex, E054 (neurospheres) ganglion eminence, E067 (brain) angular gyrus, E068 (brain) anterior caudate, E069 (brain) cingulate gyrus, E070 (brain) germinal matrix, E071 (brain) hippocampus middle, E072 (brain) inferior temporal lobe, E073 (brain) dorsolateral prefrontal cortex, E074 (brain) substantia nigra, E081 (brain) fetal brain male, E082 (brain) fetal brain female, E003 (ESC) H1 cells, E008 (ESC) H9 cells, E007 (ES-derived) H1 derived neuronal progenitor cultured cells, E009 (ES-derived) H9 derived neuronal progenitor cultured cells and E010 (ES-derived) H9 derived neuron cultured cells. A FDR of 1×10^−6^ was applied to identify significant interactions (default parameter), separately for each analysis.

### Gene-based association analysis

Genome-wide gene-based association analysis was performed using mvGWAS summary statistics as input into MAGMA (v1.07)^41^, using default parameters implemented in FUMA (SNP-wide mean model). This process examines the joint association signals of all SNPs within a given gene (including 50kb upstream to 50kb downstream of the gene), while considering the LD between the SNPs. SNPs were mapped to 20,146 protein-coding genes based on NCBI 37.3 gene definitions, and each gene was represented by at least one SNP. We applied a Bonferroni correction for the number of tested genes (p<0.05/20,146).

### Gene-set enrichment analysis

We used MAGMA^41^, again with default settings as implemented in FUMA, to test for enrichment of association within predefined gene sets. This process tests whether gene-based P values among all 20,146 genes are lower for those genes within pre-defined functional sets than the rest of the genes in the genome, while correcting for other gene properties such as the number of SNPs. A total of 7,343 gene sets, defined according to Gene Ontology biological processes, were tested from MSigDB version 7.0^45^. In the main text we report the gene-sets with p-values that met Bonferroni correction for multiple testing (p<0.05/7,343).

In addition, we used the list of single closest genes to the 27 lead SNPs arising from mvGWAS (Table 1) as input for gene set enrichment analysis, using the same 7,343 GO biological process gene sets, but now based on the hypergeometric test as implemented in GENE2FUNC of FUMA^17^, which is appropriate for gene lists.

### Protein-protein interaction network

We used the Search Tool for the Retrieval of Interacting Genes/Proteins (STRING; http://string-db.org)^44^ for protein network analysis, using as input the names of 62 overlapping genes between those identified from SNP-to-gene mapping and those from gene-based association analysis, as described above. The STRING dataset includes protein-protein interaction information from numerous sources, including experimental data, publications and computational prediction methods. Only links with medium confidence or higher (confidence score>0.4; default parameter) were retained.

### Developmental stage analysis

Using the gene-based association p-values for all 20,146 genes genome-wide, we used MAGMA (default settings as implemented in FUMA) to examine whether genetic association with brain asymmetry in our data was related to differential mRNA expression (significantly higher gene expression) in BrainSpan^47^ data from any particular ages, separately for 29 different age groups ranging from 8 postconceptional weeks to 40 years old, and 11 defined developmental stages from early prenatal to middle adulthood. We corrected for multiple testing through a FDR of 0.05 (separately for the two analyses).

### Genetic overlap of brain asymmetry with brain disorders, behavioral and cognitive traits

We applied the iSECA^53^ toolbox that can test for genetic overlap based on per-SNP association p values only (as mvGWAS does not produce univariate beta coefficient effect size estimates that can be used in standard genetic correlation analysis). We tested for genetic overlap in relation to traits previously reported to associate phenotypically with different aspects of brain structural asymmetry (see main text), using GWAS P values from previously published, large-scale studies: educational attainment (n=1,131,881)^42^, handedness (n=331,037)^22^, intelligence (n=269,867)^43^, autism spectrum disorder (n=46,350)^60^, attention deficit hyperactivity disorder (n=55,374)^55^, and schizophrenia (n=150,064)^73^. We also tested for genetic overlap in relation to brain intracranial volume (n=32,438)^78^. After LD-based filtering and clumping using default parameters, iSECA tests for pleiotropy between two sets of GWAS results using an exact binomial statistical test at each of 12 p-value levels: p≤(0.01, 0.05, 0.1, 0.2, 0.3, 0.4, 0.5, 0.6, 0.7, 0.8, 0.9, 1). The analysis compares the expected and observed overlap in the subsets of SNPs at these levels from two GWAS (144 combinations in total). In other words, iSECA iterates through each of the 12 p-value levels and counts the number of overlapping variants between two GWAS at each p-value threshold, and compares that number to the number expected under the null hypothesis of no genetic overlap, using the exact binomial test. iSECA then counts up the number of comparisons with evidence of overlap at a nominally significant level of p≤0.05. To assess the significance level of overlap, we generated 1000 data sets through permutations (default parameter), which contained all the possible combinations for a pair of traits, and determined if the number of levels with nominally significant genetic overlap was significantly more than expected by chance. Finally, Bonferroni correction <0.05 was applied for multiple testing of seven traits. Additionally, iSECA generated Q-Q plots for asymmetry mvGWAS p-values conditioned on the other trait p-values (e.g. p≤0.1, 0.2, 0.3, 0.4, 0.5, 0.75, 1.0) to visualize whether there is an excess of pleiotropic SNPs, which should be visible as a leftward shift of the curve as the P-value threshold becomes tighter (Supplementary Fig. 8).

## Acknowledgements

The research was funded by the Max Planck Society (Germany) and grants from the Netherlands Organization for Scientific Research (NWO) (054-15-101) and French National Research Agency (ANR, grant No. 15-HBPR-0001-03) as part of the FLAG-ERA consortium project ‘MULTI-LATERAL’, a Partner Project to the European Union’s Flagship Human Brain Project. Many thanks to Nathalie Tzourio-Mazoyer and Antonietta Pepe for contributions to the MULTI-LATERAL project. This research was conducted using the UK Biobank resource under Application Number 16066, with Clyde Francks as the principal applicant. Our study made use of imaging-derived phenotypes generated by an image-processing pipeline developed and run on behalf of UK Biobank^90^.

## Data Availability

The primary data used in this study are available via the UK Biobank website www.ukbiobank.ac.uk. Other publicly available data sources and applications are cited in the Methods section.

## Code availability

All code used for these described analyses is available upon request from the author.

## Competing interests

The authors declare that they have no competing interests.

## Author contributions

Z.S.: Conceptualization, methodology, analysis, visualization, original draft writing, review & editing. D.S.: Methodology, analysis, bioinformatics, review & editing. A.C-C.: Conceptualization, methodology, analysis, visualization, review & editing. M.J. Conceptualization, supervision, funding acquisition, review & editing. B. M.: Conceptualization, supervision, funding acquisition, review & editing. S. E. F.: Conceptualization, funding acquisition, review & editing. F. C.: Conceptualization, supervision, original draft writing, funding acquisition, review & editing. C. F.: Conceptualization, direction, funding acquisition, supervision, original draft writing, review & editing.

## Notes

### Competing Interest Statement

The authors have declared no competing interest.

